# Deletion of smooth muscle O-GlcNAc transferase prevents development of atherosclerosis in western diet-fed hyperglycemic ApoE^-/-^ mice in vivo

**DOI:** 10.1101/2023.03.20.533533

**Authors:** Saugat Khanal, Neha Bhavnani, Amy Mathias, Jason Lallo, Shreya Gupta, Vahagn Ohanyan, Jessica Ferrell, Priya Raman

## Abstract

Accumulating evidence highlights protein O-GlcNAcylation as a putative pathogenic contributor of diabetic vascular complications. We previously reported that elevated protein O-GlcNAcylation correlates with increased atherosclerotic lesion formation and VSMC proliferation in response to hyperglycemia. However, the role of O-GlcNAc transferase (OGT), regulator of O-GlcNAc signaling, in evolution of diabetic atherosclerosis remains elusive. The goal of this study was to determine whether smooth muscle OGT (smOGT) plays a direct role in hyperglycemia-induced atherosclerotic lesion formation and SMC de-differentiation. Using tamoxifen-inducible *Myh11-CreER^T^*^2^ and OGT^fl/fl^ mice, we generated smOGT^WT^ and smOGT^KO^ mice, with and without ApoE-null backgrounds. Following STZ-induced hyperglycemia, smOGT^WT^ and smOGT^KO^ mice were kept on standard laboratory diet for study duration. In a parallel study, smOGT^WT^ApoE^-/-^ and smOGT^KO^ApoE^-/-^ were kept on Western diet beginning 8-wks-age. Animals harvested at 14-16-wks-age were used for plasma and tissue collection. Loss of smOGT augmented SM contractile marker expression in aortic vessels of STZ-induced hyperglycemic smOGT^KO^ mice. Consistently, smOGT deletion attenuated atherosclerotic lesion lipid burden (Oil red O), plaque area (H&E), leukocyte (CD45) and smooth muscle cell (ACTA2) abundance in Western diet-induced hyperglycemic smOGT^KO^ApoE^-/-^ mice. This was accompanied with increased SM contractile markers, and reduced inflammatory and proliferative marker expression. Further, smOGT deletion attenuated YY1 and SRF expression (transcriptional regulators of SM contractile genes) in hyperglycemic smOGT^KO^ApoE^-/-^ and smOGT^KO^ mice. These data uncover an atheroprotective outcome of smOGT loss-of-function and suggest a direct regulatory role of OGT-mediated O-GlcNAcylation in VSMC de-differentiation in hyperglycemia.

## Introduction

Cardiovascular disease (CVD) is the leading cause of morbidity and mortality world-wide, claiming about 19 million lives in 2020^1^. CVD has continued to pose tremendous economic burden on the national health care expenditure, with an estimated annual total cost of $378 billion in 2017-18. Atherosclerosis, one of the major players of cardiovascular anomalies, results in occlusion of the affected arteries triggering multiple complications including ischemic stroke, myocardial infarction, heart failure and angina^2^. Despite the well-accepted contribution of lipids to atherosclerosis, current lipid-lowering therapies including statins and PCSK9 inhibitors may offer limited benefit against reduction of major adverse cardiovascular events. Moreover, emergence of statin intolerance and associated risks of hyperglycemia coupled with the escalated costs of PCSK9 inhibitors have raised added concerns about the risk-benefit ratio of these agents^3, 4^.

Vascular disease is the most prevalent and deadliest of all health concerns affecting diabetic patients in our society. Epidemiological studies indicate that diabetic patients and individuals with impaired glucose tolerance have 2-4-fold increased risk for development of atherosclerosis compared to individuals without diabetes^5^. Multiple clinical studies and trials including animal data have shown that elevated plasma glucose levels, hallmark of diabetes, have profound proatherogenic properties, independent of hyperlipidemia^6–8^. Taken together, these findings highlight hyperglycemia as an independent risk-factor for initiation and progression of macrovascular complications. Yet, optimal glycemic control alone does not effectively lower the risks of atherosclerosis in long-standing diabetic patients^9^. Indeed, previous studies have provided evidence for the role of chronic hyperglycemia and duration of diabetes in development of atherosclerosis^10, 11^. This has led to the idea that deranged glucose metabolism following chronic hyperglycemia may be responsible for accelerated atherosclerotic complications associated with diabetes. However, the inherent molecular basis for diabetes-induced vasculopathy remains incompletely understood.

In healthy individuals, vascular smooth muscle cells (VSMC), residing in the medial layer of the vascular wall, play a key role in maintaining the normal vascular tone enabling effective flow of arterial blood throughout the body. In diabetic patients, chronic exposure to elevated circulating glucose concentrations makes these individuals highly prone to increased VSMC migration and proliferation^12^. De-differentiation of SMC into the synthetic phenotype with augmented migratory and proliferative properties accompanied with reduced contractile gene expression is a key event central to the evolution of atherosclerosis^13, 14^. Despite extensive studies, a complete molecular understanding of how hyperglycemia regulates this process is obscure.

O-GlcNAcylation is an important post-translational modification that involves the attachment of O-linked N-acetylglucosamine (O-GlcNAc) moiety to serine and threonine residues of numerous cytoplasmic, nuclear, and mitochondrial proteins^15, 16^. This process is tightly regulated by two key enzymes, O-linked N-acetylglucosaminyltransferase (OGT) and O-linked N-acetylglucosaminidase (OGA). A plethora of data indicates that O-GlcNAc protein modification is an important regulatory mechanism of intracellular glucose signaling^17^. It is widely accepted that under conditions of increased nutrient availability such as diabetes, cellular O-GlcNAc levels are profoundly elevated^18, 19^. Indeed, several studies, including our earlier work, have shown that hyperglycemia positively correlates with increased O-GlcNAcylation in various cells and tissues^17, 20, 21^. Growing literature further reinforces the notion that O-GlcNAcylation is a putative pathogenic contributor of diabetes and related vascular complications^22–26^. Previous studies have shown that incubation of vascular cells including VSMC and endothelial cells with high glucose *in vitro*, reflective of the diabetic milieu, activates the hexosamine biosynthetic pathway (HBP) resulting in sustained increase in protein O-GlcNAcylation. Augmented O-GlcNAc protein modification, in turn, mediates the upregulation of numerous genes associated with atherosclerosis including TSP-1, TGF-β, PAI-1 and NF-kB^27–30^. Earlier studies have also shown that diabetic patients may develop atherosclerotic plaques with elevated O-GlcNAc levels^31^; also, incidence of calcified plaques is profoundly enhanced in those individuals^32, 33^.

We previously demonstrated that protein O-GlcNAcylation directly associates with increased proliferation of human aortic smooth muscle cells (HASMCs) in response to high glucose in vitro^21^. In a subsequent study, we further reported that protein O-GlcNAcylation and augmented OGT expression, key controller of O-GlcNAc signaling, correlate with increased atherosclerotic lesion formation and proliferative SMC lesion abundance in hyperglycemic ApoE^-/-^ mice in vivo^34^. Recent studies have further shown that OGT, the sole enzyme mediating addition of O-GlcNAc residues to proteins, plays a crucial role in regulation of diabetic vascular calcification and diabetic wound recovery^25, 35^. However, the fundamental contribution of OGT as a driver of atherosclerotic lesion formation in diabetes has not been previously explored.

Therefore, the primary objective of the current study was to determine whether smooth muscle OGT plays a direct role in the development of hyperglycemia-induced atherosclerosis and SMC de-differentiation in diabetes. Our data demonstrates an atheroprotective consequence of SMC-specific OGT deletion in western diet-fed hyperglycemic ApoE^-/-^ mice in vivo.

## Materials and Methods

### Mouse models

All animal procedures including mice euthanasia were conducted according to protocols reviewed and annually approved by the Institutional Animal Care and Use Committee at Northeast Ohio Medical University, in accordance with the NIH guidelines for the Care and Use of Laboratory Animals. SMC-specific *Ogt* knockout mice (smOGT^KO^) were generated by crossing female homozygous OGT-floxed (OGT^fl/fl^) mice (B6.129-*Ogt^tm1Gwh^*/J; JAX # 004860) with male tamoxifen (tmx)-inducible hemizygous *Myh11-CreER^T^*^2^ mice (Cre^tg^; JAX # 019079) purchased from The Jackson Laboratories (Bar Harbor, ME). The OGT^fl/fl^ mice have loxP restriction sites on either side of the exon encoding amino acids 206-232 of the x-linked *Ogt* gene. The *Myh11-CreER^T^*^2^ mice express CreER^T2^ under the control of the mouse smooth muscle myosin heavy polypeptide 11 (*Myh11*, aka *sm-mhc*) promoter. The resulting OGT^fl/Y^/Cre^tg^ male mice (produced in F1 generation) were initially used for Cre recombinase activation to test for the successful generation of SMC-specific knockouts. **Please note**, *Ogt* floxed gene being x-linked, all male mice produced were OGT^fl/Y^ and females were OGT^fl/fl^. Also, given that Cre transgene is Y-linked, only male mice carried the Cre transgene. To generate the experimental mice for this study, male OGT^fl/Y^/Cre^tg^ mice were bred with female OGT^fl/+^; the resulting OGT^fl/Y^/Cre^tg^ and age-matched OGT^+/Y^/Cre^tg^ littermate mice were used for Cre recombinase activation. In a concurrent study, OGT^fl/Y^/Cre^tg^ male mice were cross-bred with ApoE^-/-^ female mice (JAX stock #002052) to generate Cre^tg^/OGT^fl/Y^/ApoE^-/-^ and age-matched Cre^tg^/OGT^+/Y^/ApoE^-/-^ littermates subsequently utilized for Cre recombination. OGT^F^, Cre^tg^ and ApoE^-/-^ genotypes were confirmed by polymerase chain reaction, per modification of established protocols (The Jackson Laboratory). All mice utilized in this study were on C57BL6/J background, housed in a pathogen-free environment and maintained on 12:12 h light/dark cycle.

### Study Design

In the first study, mice genotypes weaned at 4-wks-age were maintained on standard laboratory diet (Purina LabDiet 5008) provided *ad libitum* until study end point. Briefly, upon weaning mice were randomly allocated to the following groups: i) tamoxifen-treated Cre^tg^/OGT^fl/Y^ (smOGT^KO^) ii) vehicle-treated Cre^tg^/OGT^fl/Y^ (smOGT^WT^) and iii) tamoxifen-treated Cre^tg^/OGT^+/Y^ (smOGT^WT^). For Cre recombinase activation, 6-wks-old male mice were treated with tamoxifen (tmx, 60 mg/kg/day) or vehicle (corn oil) intraperitoneally once daily for 5 consecutive days. After allowing for one-week ‘washout’, mice were injected with either low-dose STZ (50mg/kg/day) or sodium citrate buffer, pH7.4 (vehicle control) intraperitoneally once daily for five consecutive days beginning at 8-wks-age for the induction of hyperglycemia. Ten days after initiation of STZ, blood samples collected by lateral tail incision were used for glucose estimation using a one-touch glucometer. Mice with non-fasted blood glucose levels ≥ 250 mg/dl were identified as hyperglycemic and assigned to the study for comparison with the normoglycemic genotypes. In a parallel study, 6-wks-old Cre^tg^/OGT^fl/Y^/ApoE^-/-^ and age-matched Cre^tg^/OGT^+/Y^/ApoE^-/-^ littermate male mice were subjected to tmx-induced Cre recombination as described above, resulting in development of smOGT^KO^ and smOGT^WT^ mice on ApoE^-/-^ backgrounds, respectively. At 8-wks-age, mice genotypes were subjected to Western Diet (TD88137, Teklad, Harlan) feeding regimen. In each study, body weight and non-fasted blood glucose levels were monitored every two weeks in all animals. After an overnight fasting, mice were harvested at 14-16 weeks of age for blood and tissue collection following euthanasia using Fatal Plus.

### Plasma Lipid Analyses

Plasma separated from the blood samples collected from the mice genotypes were utilized for estimation of plasma total cholesterol and total triglyceride levels using Infinity Reagents (ThermoFisher).

### Glucose Tolerance Test

Following overnight fasting, a lateral tail incision was made to obtain an initial blood sample to measure the baseline (fasting) plasma glucose in each mouse. This was followed by a single intraperitoneal injection of sterile glucose solution (2 g/Kg body weight) to conduct Glucose Tolerance Test (GTT). Blood samples were periodically collected 15, 30, 60, 90, and 120 minutes after the glucose injection. For each time point, a hand-held glucometer was utilized for the measurement of blood glucose levels.

### Echocardiographic Analysis of Cardiac Function

At 16-wks-age prior to harvest, cardiac function was measured in each mouse via echocardiography using the Vevo 770 system (VisualSonics), equipped with a 710B-075 transducer (20–30 MHz) specifically designed for small animal investigations at a frame rate of 40–60 Hz, as described earlier^36^. Images were collected in M-mode from the parasternal short axis view at mid-papillary muscle level. Briefly, each mouse was placed on a controlled heating platform made for small animal echocardiography following anesthesia. A nose cone was utilized to deliver 1-2% isoflurane at 0.5 liters/min with oxygen for induction of anesthesia. Following removal of chest hair, Aquasonic 100 gel (Parker Laboratories) pre-warmed to 37°C was applied to the chest followed by assessment of cardiac function. All echocardiographic measurements and calculations were performed offline using the Vevo770/3.0.0 software. Five cycles were used to average all measurements.

### Metabolic Phenotyping

Prior to harvest, a subset of mice genotypes was subjected to metabolic phenotyping using the Comprehensive Lab Animal Monitoring System (CLAMS). Briefly, mice were kept individually in sealed Plexiglass cages through which fresh room air was provided at 0.5 liters/minute for the entire duration of the experiment which included 24 hours of acclimatization followed by at least 24 hours of normal recording. During this time, the animals had unlimited access to food and water and indirect calorimetry was used to assess their metabolic performance. For each mouse, oxygen consumption (VO_2_), carbon dioxide production (VCO_2_), respiratory exchange ratio (RER [VCO_2_/VO_2_]), and energy production was measured and recorded over the course of 24 hours. Furthermore, an EchoMRI 3-in-1 body composition Analyzer was used to measure both fat and lean tissue mass in live mice prior to CLAMS.

### Aortic Root Morphometry

At study endpoint, mouse hearts were removed after PBS perfusion. After a brief PBS wash, the individual mouse hearts were OCT-embedded and stored at −80°C for further processing. Following dissection of the OCT-embedded hearts, serial sections (8 microns) of the aortic root were obtained, as reported earlier^37^. For all morphometric measurements, serial sections within 100-150 microns from the valve leaflet were utilized. Moreover, care was taken to ensure that only sections from comparable areas of the aortic root across all treatment groups were included for further staining and quantification. Concurrent aortic root sections derived from each mouse were stained with 0.5% Hematoxylin and Eosin (H & E) and Oil red O (ORO) solutions to detect lipid burden and plaque area, respectively. Further, hematoxylin was used as a counterstain for all ORO-stained sections. Sections were mounted using the DPX mounting media and images were captured at 10X magnification using the Olympus BX61VS microscope. For quantitative morphometry, we utilized a total of 7 mice for each genotype, and at least 30 sections from each group were analyzed.

### Immunohistochemistry

Serial aortic root sections derived from each animal were subjected to immunohistochemistry using anti-CD45 (Bioss, Woburn, MA) and anti-αSMA (Sigma, St. Louis, MO) antibodies. Briefly, tissue sections were incubated in ice-cold acetone for 5–10 minutes followed by blocking with 5% donkey or goat serum at room temperature for an hour. After an overnight incubation at 4°C with primary antibodies (anti-αSMA-1:200; anti-CD45-1:200) diluted in the blocking solution, sections were incubated with Alexa Flour 594 donkey anti-rabbit IgG secondary antibody (1:400, Vectashield, Vector Laboratories) at room temperature for 1 hour. To confirm non-specific staining, identical root sections were concurrently incubated either in the absence of the corresponding primary antibody or with species-specific isotype-matched IgG control antibody relevant to the primary antibody. Images were acquired using the Olympus fluorescence IX71 microscope at 10X magnification with an identical set of parameters applied across all sections, specific for each antibody.

### Immunoblotting and Quantitative real time PCR

Aortic vessels isolated from the mice genotypes were utilized in immunoblotting. Briefly, aortic protein lysates were prepared in RIPA lysis buffer and protein concentration was measured using the BCA Gold protein assay (ThermoFisher). Equal protein amounts (25μg) of each sample were resolved on 8% SDS-PAGE followed by a wet transfer to PVDF membranes. Immunoblotting was performed using the following antibodies: OGT, α-SMA, YY1, SRF, pERK and tERK (1:1000, Cell Signaling, Danvers, MA); O-GlcNAc (RL2, 1:1000) and PCNA (1:500, Abcam); LMOD1 (1:1000, Proteintech). Equal protein loading was confirmed by staining the membranes with Ponceau S, used as a loading control. All immunoblot images were captured using Protein Simple and densitometric analyses were performed using the Image J software. For mRNA analysis, total RNA was extracted from the aortic vessels isolated from each mouse using Trizol Reagent (Sigma Aldrich, St. Louis, MO), per manufacturer’s instructions. RNA concentrations were determined using a nanodrop spectrophotometer (ThermoFisher, Waltham, MA). Two micrograms of total RNA were converted into cDNA using the RETROscript Reverse Transcription Kit (ThermoFisher). Quantitative RT-PCR was performed using 2μl of the reverse transcribed cDNA, relevant forward and reverse primers and 1X SYBR PCR master mix in CFX96 PCR system (Bio-Rad Laboratories). mRNA expression levels were normalized using the housekeeping genes 36b4 and 18S. All RT and PCR reactions were conducted in duplicate for each sample with and without RT as controls; cycle threshold (C_t_) values were converted to relative gene expression levels using the 2-^ΔΔC(t)^ method.

### Statistical Analysis

Statistical analyses were performed using GraphPad Prism version 8. Each data set was tested for normality (Shapiro-Wilks; Kolmogorov-Smirnov) and homoscedasticity (Brown-Forsythe) prior to further analysis. When the assumption of normality was met, parametric testing methods were used whereas non-parametric methods were employed when the conditions of normality were violated. Ordinary one-way ANOVA was used for normally distributed data with similar variance followed by Tukey HSD post-hoc test. For normally distributed data with unequal variance, Welch ANOVA was applied followed by Games-Howell post-hoc test. Mann-Whitney and Kruskal-Wallis were used as the non-parametric counterparts of student’s t-test and one-way ANOVA, respectively; Kruskal-Wallis was followed by the Dunn’s post-hoc test. Statistical significance was considered at p ≤ 0.05. Each experiment was repeated at least three times in an independent setting, with two to five replicates for each treatment. In immunofluorescence experiments, six to eight independent field pictures were acquired for each individual treatment in a single experiment. For aortic root morphometry and immunohistochemistry, four to five sections per mouse and three to eight mice per group were examined. For all immunoblots, lane images depict proteins loaded and detected on a single blot. However, for specific immunoblots as indicated in the corresponding figure legends, lanes were rearranged for the purpose of clarity. All data is presented as fold-increase vs. controls. Values are expressed as Mean ± SEM.

## Results

### Validation of inducible SMC-specific OGT knockout mice

To study the role of smooth muscle OGT, we generated inducible SMC-specific *Ogt* knockout mice (**Figure 1A**) by crossing female homozygous OGT floxed mice with tmx-inducible *Myh11-CreER^T^*^2^ (hemizygous Cre^tg^) male mice, expressing CreER^T2^ under control of the mouse smooth muscle myosin heavy chain 11 promoter. **Figure 1B** shows genotyping data confirming generation of OGT^fl/Y^/Cre^tg^ and OGT^+/Y^/Cre^tg^ mice. Mice were injected with Tmx or corn oil as described in Methods for Cre recombinase activation. This was followed by collection of the aortae and left ventricular tissue at least 3 weeks post-Tmx. Immunoblotting confirmed significant reduction in OGT expression that was noted only in aortic lysates prepared from Tmx-treated OGT^fl/Y^/Cre^tg^ mice. Specifically, OGT expression was reduced by about 70% in aortic vessels derived from Tmx-treated OGT^fl/Y^/Cre^tg^ mice vs. from Tmx-treated OGT^+/Y^/Cre^tg^ littermates (**Figure 1C**, **left panel**; p<0.0001). In contrast, OGT expression was unaffected in the left ventricle (LV) tissue lysates prepared from these mice genotypes under similar experimental conditions. These results validated successful generation of inducible SMC-specific OGT knockout murine model (smOGT^KO^).

**Figure 1.**
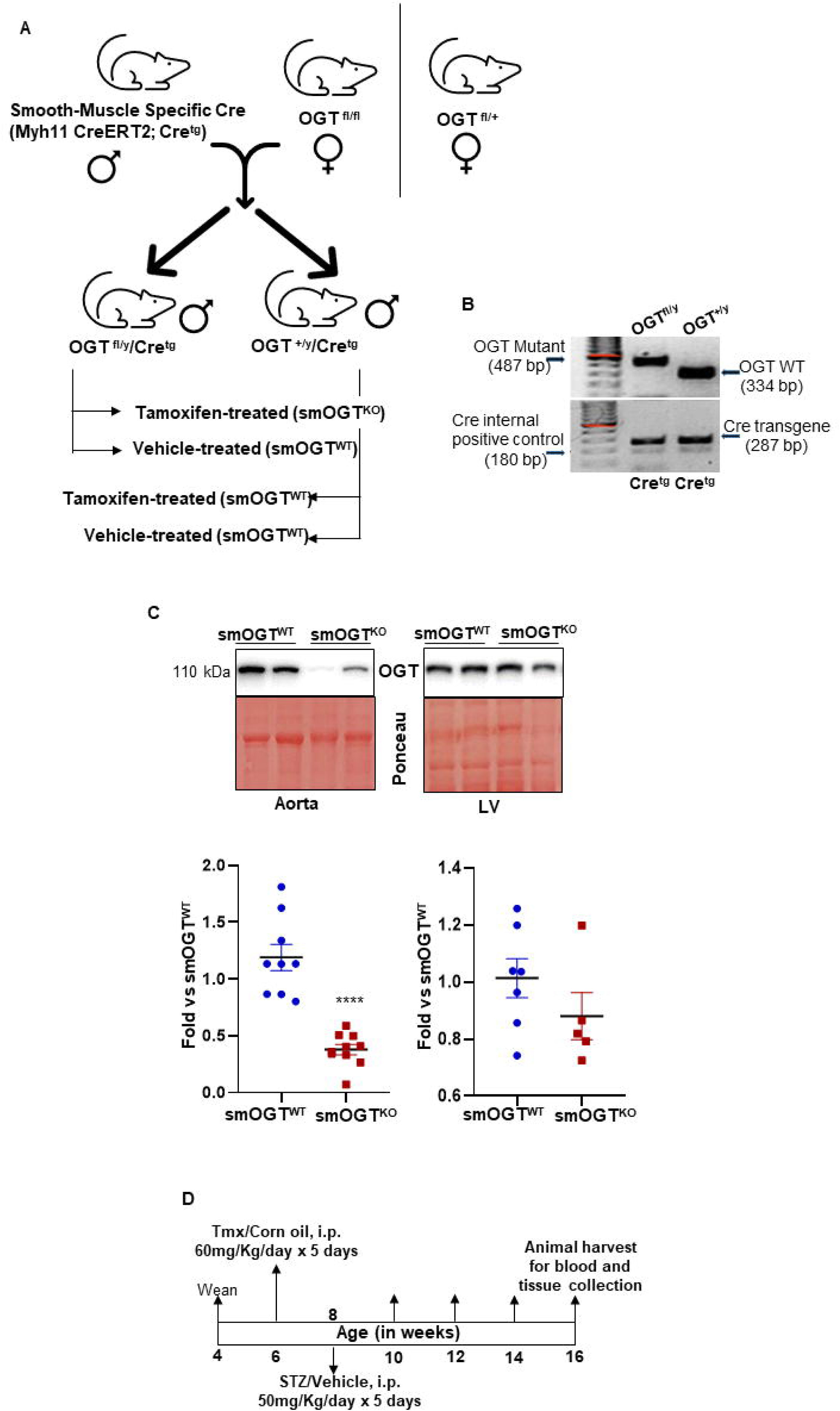
Validation of inducible SMC-specific OGT knockout mice. A) Breeding strategy, B) genotyping data, C) representative immunoblots and summary graphs showing OGT expression in aorta vs left ventricle isolated from smOGT^KO^ vs smOGT^WT^ mice, D) study time line for STZ-treated hyperglycemic smOGT^WT^ vs smOGT^KO^ mice.

### Metabolic phenotype of SMC-specific OGT knockout mice under basal and STZ-induced hyperglycemia

To induce hyperglycemia reflective of diabetes, smOGT^WT^ and age-matched smOGT^KO^ mice were treated with STZ or sodium citrate (vehicle control) intraperitoneally, as described in Methods. Body weight and random blood glucose levels were monitored every two weeks in all animals until harvest at 16-weeks-age (**Figure 1D**). As shown in **Figure 2A**, no significant differences in body weights were observed in smOGT^KO^ vs. smOGT^WT^ mice treated with or without STZ. Random blood glucose monitoring revealed significantly elevated glucose levels (>250mg/dl) in STZ-treated mice genotypes, as early as 10 days following initiation of STZ administration. Intraperitoneal glucose tolerance test (IPGTT) was conducted to examine glucose responsiveness in the mice genotypes. As expected, STZ administration significantly lowered the ability of the mice to handle an intraperitoneal glucose load and this effect was observed in both smOGT^WT^ and smOGT^KO^ mice (**Figure 2B**). Moreover, no statistically significant difference in plasma total cholesterol and total triglyceride levels were noted between smOGT^WT^ and smOGT^KO^ mice under STZ- and non-STZ treated conditions (**Figures 2C, 2D**). In addition, echocardiographic measurements revealed unaltered Ejection Fraction (%EF), Fractional Shortening (%FS), left ventricular internal diameter (LVID) and left ventricular volume (LV vol) in smOGT^WT^ vs. smOGT^KO^ mice, and this effect was consistent across both non-hyperglycemic and STZ-induced hyperglycemic mice (**Figures 2E-2H**). Indirect calorimetry using CLAMS demonstrated no significant difference in oxygen consumption (VO_2_, **Figure 3A**), carbon dioxide production (VCO_2_, **Figure 3B**), RER, a measure of fuel usage (**Figure 3C**) and other metabolic parameters including physical activity (**Figure 3D**), energy expenditure (**Figure 3E**) and food intake (**Figure 3F**) between non-hyperglycemic and hyperglycemic mice with and without SMC-specific OGT deletion. Further, assessment of fat and lean mass using Echo MRI did not show any difference in adiposity between any of the mice treatment groups (**Figure 3G**). Together, these results clearly demonstrate that SMC-specific loss of OGT does not adversely affect the metabolic phenotype and cardiac function of STZ-induced hyperglycemic mice *in vivo*.

**Figure 2.**
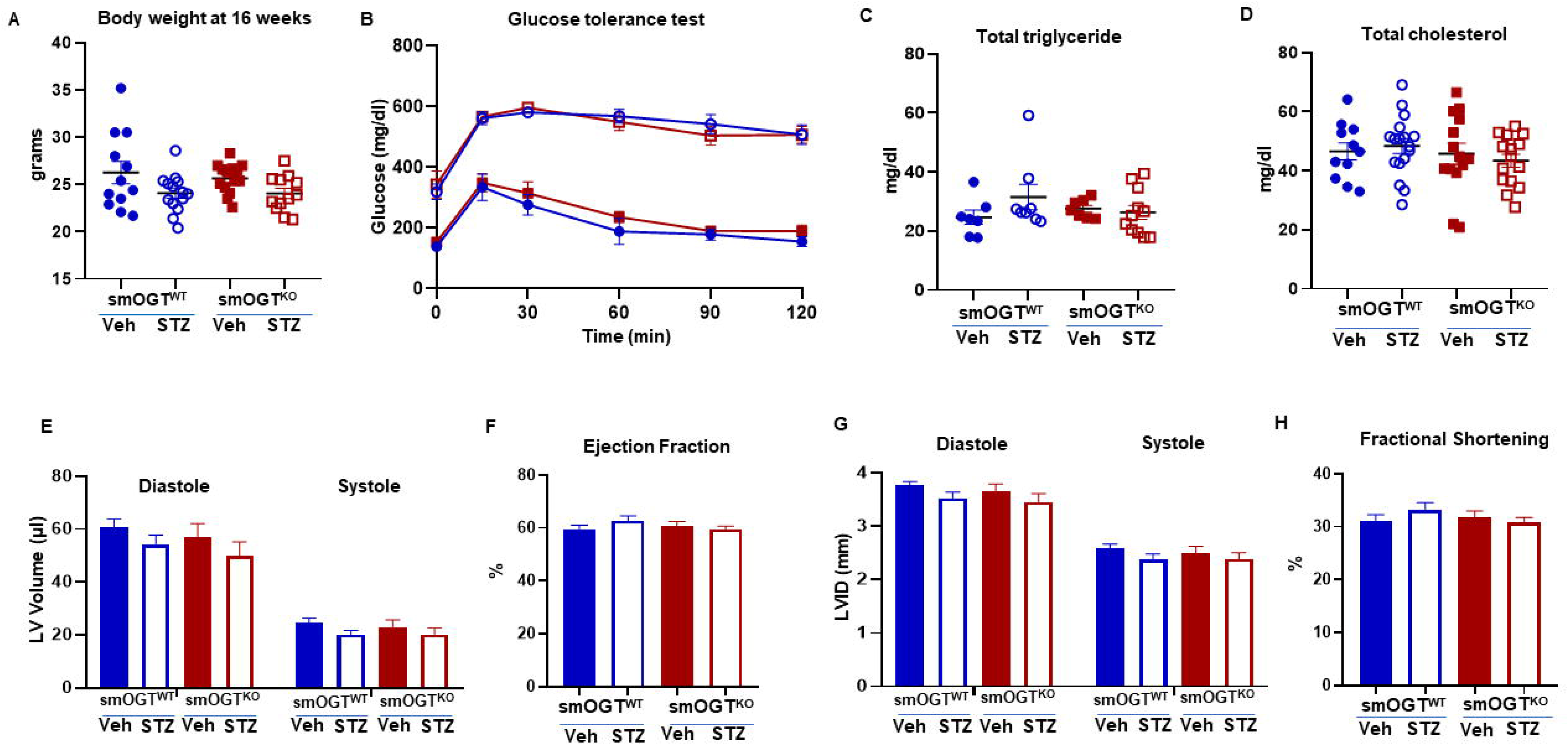
Metabolic and cardiac phenotypes of SMC-specific OGT knockout mice under basal and STZ-induced hyperglycemic conditions. Male smOGT^WT^ and smOGT^KO^ mice were treated with STZ or sodium citrate (vehicle control) as shown in the study timeline in figure legend 1. Shown are A) body weight, B) Glucose tolerance test, C) Total triglyceride, D) total cholesterol, E) LV volume, F) ejection fraction, G) LV internal diameter (LVID) and H) fractional shortening of vehicle- and STZ-treated smOGT^WT^ and smOGT^KO^ mice at 16-wks-age.

**Figure 3.**
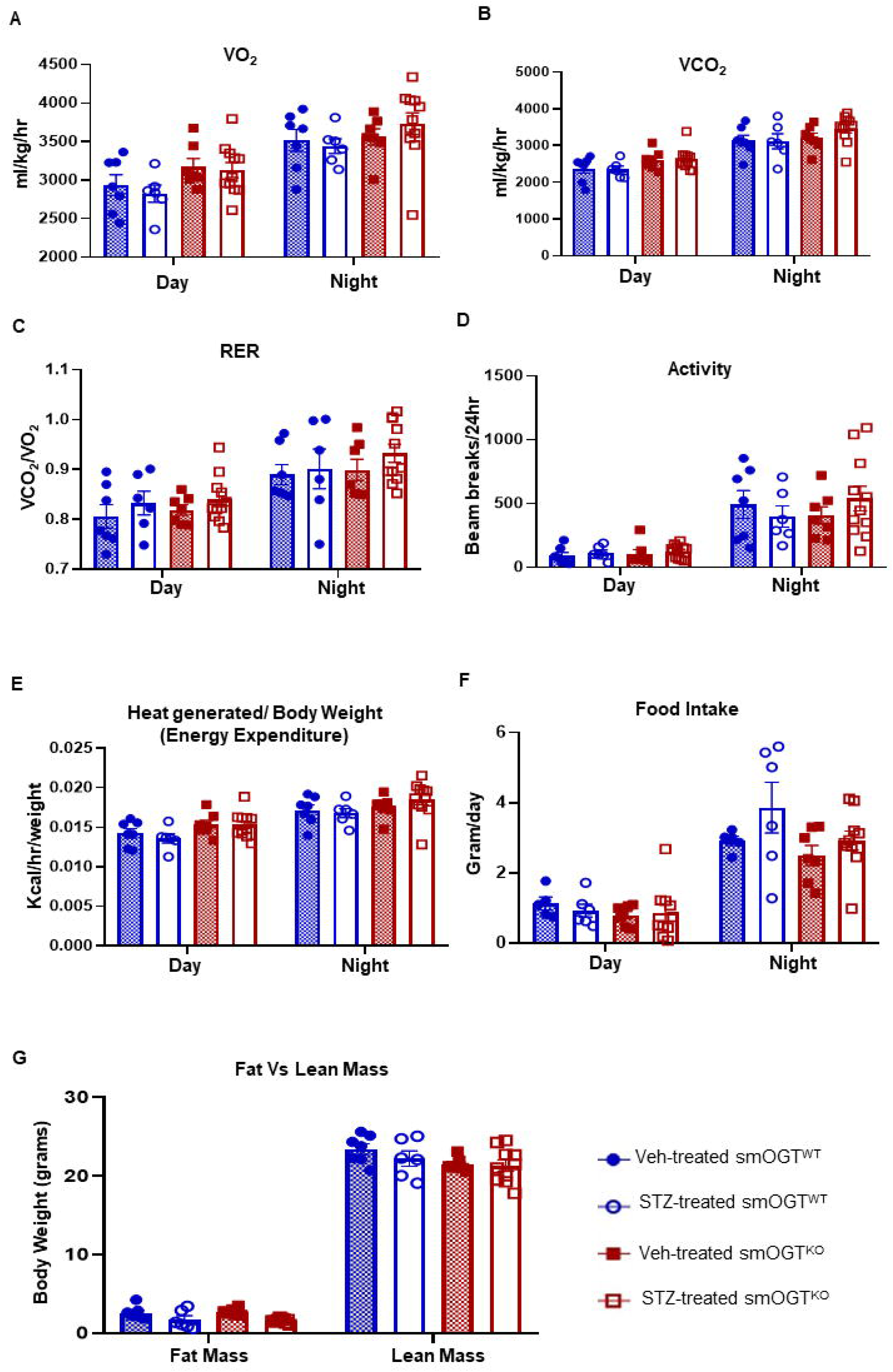
Indirect calorimetry of SMC-specific OGT knockout mice under basal and STZ-induced hyperglycemia via CLAMS.

### Reduced O-GlcNAc protein expression increases SM contractile marker expression in the aortic vasculature of STZ-induced hyperglycemic mice with SMC-specific OGT deficiency

To determine whether smooth muscle OGT plays a direct role in VSMC differentiation in diabetes, aortic lysates derived from non-hyperglycemic and STZ-induced hyperglycemic smOGT^WT^ and smOGT^KO^ mice were subjected to immunoblotting, as described in Methods. As shown in **Figures 4A and 4B**, OGT and O-GlcNAc protein expression were upregulated in the aortic vessels of smOGT^WT^ mice following STZ-induced hyperglycemia. In contrast, SMC-specific OGT deletion significantly reduced OGT and O-GlcNAc protein levels in the aortic vasculature of smOGT^KO^ vs. smOGT^WT^ mice under non-hyperglycemic and hyperglycemic conditions. Specifically, while STZ-induced hyperglycemia increased OGT and O-GlcNAc protein expression in smOGT^WT^ mice (>1.5-fold vs. non-hyperglycemic smOGT^WT^), SMC-specific lack of OGT resulted in a signifcant decrease in OGT expression and protein O-GlcNAcylation in smOGT^KO^ mice (p<0.0001 compared to smOGT^WT^), and this effect was observed both basally and under hyperglycemic conditions. Importantly, SMC-specific OGT loss and reduced protein O-GlcNAcylation increased SM contractile marker expression. Specifically, in aortic vessels of hyperglycemic smOGT^KO^ mice, ACTA2 expression was markedly increased compared to hyperglycemic smOGT^WT^ mice (p<0.0001, **Figure 4C**). Likewise, SMC-specific lack of OGT augmented LMOD1 expression in both non-hyperglycemic and hyperglycemic smOGT^KO^ mice vs. animals with intact OGT (p<0.005, **Figure 4D**). Further, real-time PCR demonstrated a statistically significant increase in *Myh11* mRNA expression in smOGT^KO^ vs. smOGT^WT^ mice in response to STZ-induced hyperglycemia (p<0.05; **Figure 4E**). Collectively, these data clearly demonstrate a direct regulatory role of smooth muscle OGT on VSMC de-differentiation in hyperglycemia, reflective of diabetes.

**Figure 4.**
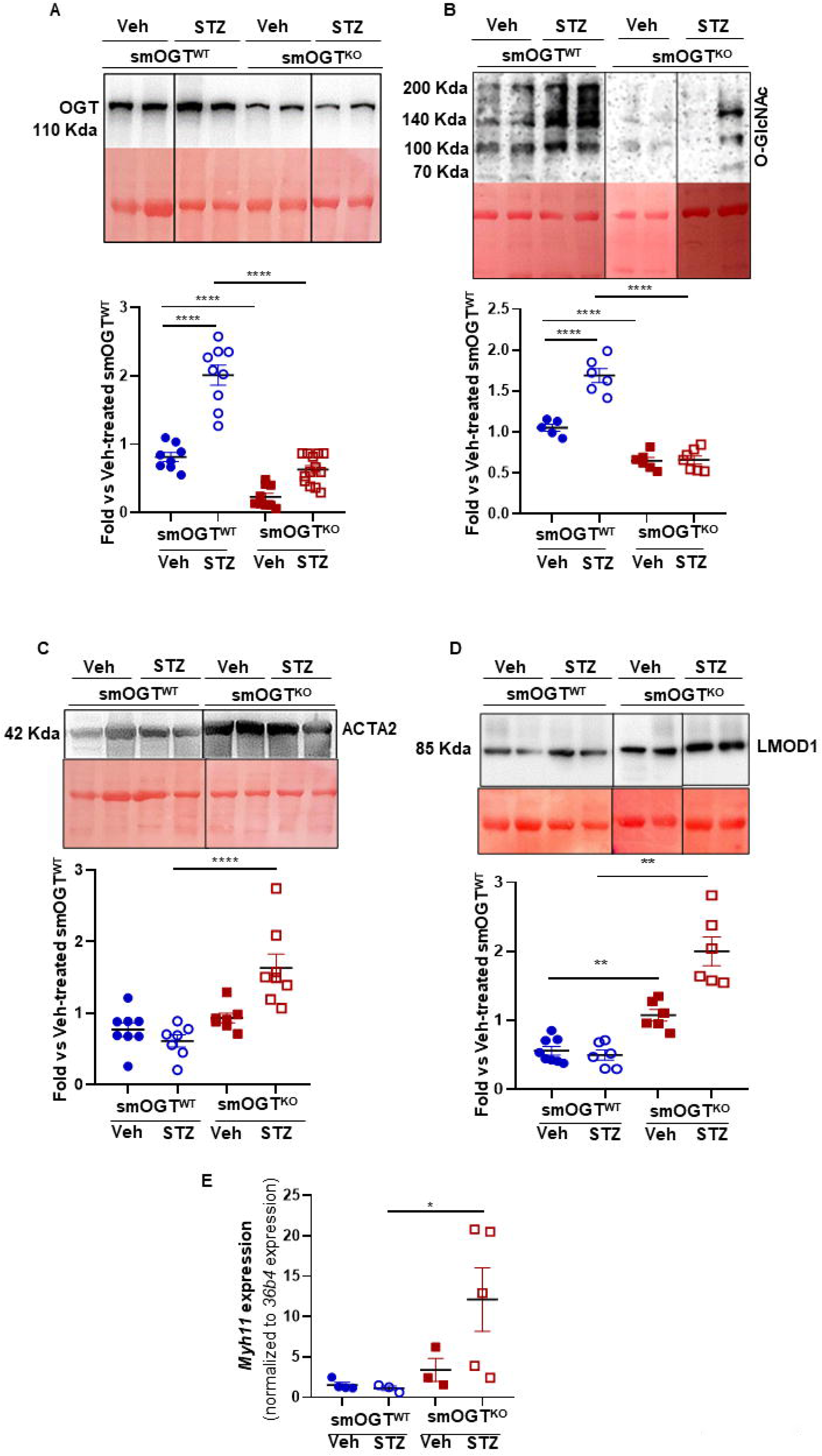
SMC-specific OGT deletion increases SM contractile marker expression in aortic vessels of STZ-induced hyperglycemic mice in vivo. A-D) immunoblotting of aortic lysates derived from vehicle- and STZ-treated smOGT^WT^ and smOGT^KO^ mice. Shown are representative immunoblots (upper panels) and corresponding summary graphs (lower panels) ^-/-^for A) OGT, B) O-GlcNAc, C) ACTA2 and D) LMOD1. E) relative mRNA expression of *Myh11*.

### Effect of SMC-specific OGT deletion on glycemic index and plasma lipid levels in western diet fed-ApoE^-/-^ mice *in vivo*

To interrogate whether smooth muscle OGT plays a direct role in development of atherosclerosis, we next generated smOGT^KO^ mice on ApoE^-/-^ atherosclerotic background. Immunoblotting of aortic lysates revealed a significant decrease in OGT and O-GlcNAc protein expression in tmx-treated Cre^tg^/OGT^fl/Y^/ApoE^-/-^ mice compared with age-matched tmx-injected Cre^tg^/OGT^+/Y^/ApoE^-/-^ littermate mice (**Figure 5A-5C**), validating the smOGT^KO^/ApoE^-/-^ murine model. Mice genotypes were subjected to a Western diet feeding regimen, as illustrated in **Figure 6A**. Western diet feeding significantly increased the fasting blood glucose levels (p<0.0001) in smOGT^WT^ApoE^-/-^ mice vs. regular chow-fed smOGT^WT^ApoE^-/-^ mice (**Figure 5D**), reflective of hyperglycemia characteristic of diabetes. Notably, hyperglycemia was accompanied with a significant elevation in O-GlcNAc protein expression in aortic vessels isolated from the Western diet-fed smOGT^WT^ApoE^-/-^ vs. standard chow-fed smOGT^WT^ApoE^-/-^ mice (**Figure 5E**). Interestingly, under similar experimental conditions, SMC-specific OGT ablation had no effect on the body weight, fasting blood glucose, glucose tolerance and plasma lipid levels in western diet-fed hyperglycemic smOGT^KO^/ApoE^-/-^ compared with hyperglycemic smOGT^WT^/ApoE^-/-^ mice with intact OGT (**Figure 6B-6G**). These results demonstrate that SMC-specific lack of OGT does not affect the glycemic index of Western diet-fed ApoE^-/-^ *in vivo*.

**Figure 5.**
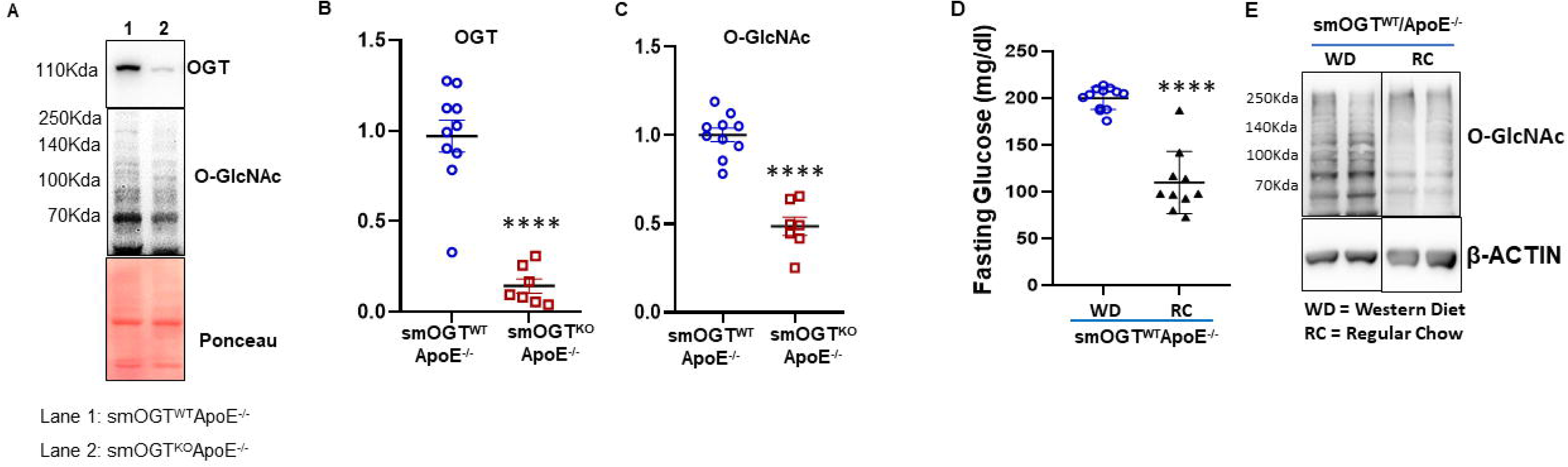
A-C) Validation of SMC-specific OGT deletion in ApoE^-/-^ mice. D-E) Western diet feeding increases fasting blood glucose levels accompanied with increased O-GlcNAc protein expression in the aortic vessels of ApoE^-/-^ mice.

**Figure 6.**
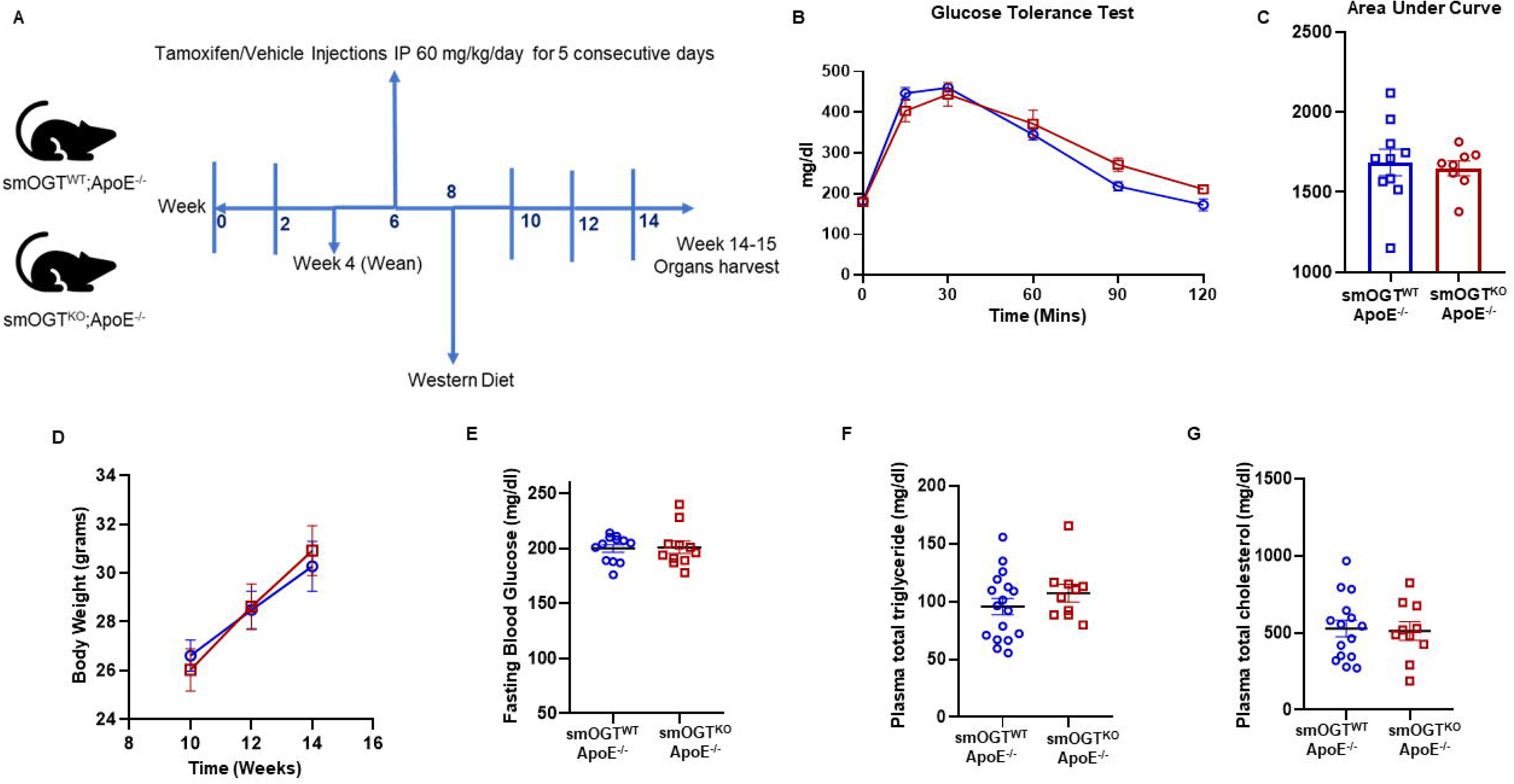
Metabolic profile of western diet-fed ApoE^-/-^ mice with SMC-specific OGT deletion. Male smOGT^WT^;ApoE^-/-^ and smOGT^KO^;ApoE^-/-^ were subjected to Western diet feeding for 6 wks. A) study timeline, B) glucose tolerance test (GTT), C) GTT Area under Curve, D) body weight, E) fasting blood glucose, F) total triglyceride and G) total cholesterol.

### SMC-specific OGT deficiency impedes lesion burden in western diet-fed hyperglycemic ApoE^-/-^ mice

Using aortic root morphometry, we analyzed atherosclerotic lesion formation in hyperglycemic smOGT^WT^ApoE^-/-^ and smOGT^KO^ApoE^-/-^ mice following 6-7 weeks of Western diet feeding. As shown in **Figure 7A**, Oil red O (ORO) staining of aortic root sections revealed large lipid-filled lesions in smOGT^WT^ApoE^-/-^ mice. In contrast, SMC-specific OGT deletion significantly attenuated the lipid burden in smOGT^KO^ApoE^-/-^ aortic roots (**Figure 7A**). Quantification of the ORO-positive area demonstrated a statistically significant reduction (p<0.0001) in lipid-filled lesions that were formed in the aortic sinus of smOGT^KO^ApoE^-/-^ mice vs. smOGT^WT^ApoE^-/-^ with intact OGT, in response to Western diet-induced hyperglycemia (**Figure 7C**). To assess the plaque area in the mice genotypes, we next performed H & E staining of the aortic root sections derived from our mice cohorts. As shown in **Figure 7B**, hyperglycemic smOGT^WT^ApoE^-/-^ mice with intact OGT showed remarkable plaque area that was consistent with the increased lipid burden in these animals as compared with hyperglycemic ApoE^-/-^ mice lacking SMC-specific OGT. Specifically, quantification of H & E staining revealed marked decrease in aortic root lesion area in smOGT^KO^ApoE^-/-^ vs. smOGT^WT^ApoE^-/-^ with intact OGT following western diet feeding (p<0.0001; **Figure 7D**). Taken together, these data clearly demonstrate an atheroprotective effect of SMC-specific OGT deficiency in Western Diet-fed hyperglycemic ApoE^-/-^ mice *in vivo*.

**Figure 7.**
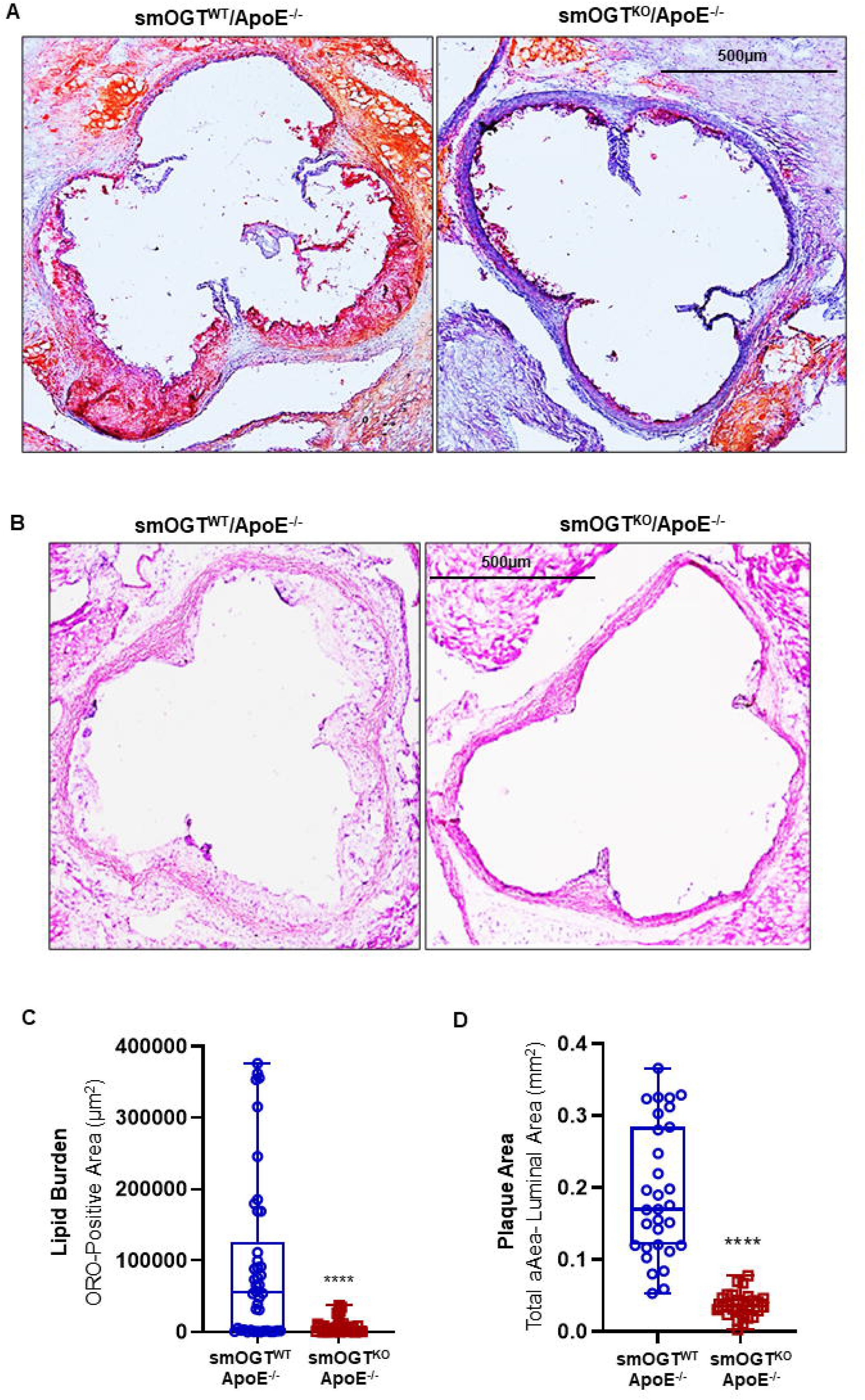
SMC-specific OGT deletion impedes lipid burden and plaque area in Western diet-fed ApoE^-/-^ mice. Shown are representative A) ORO- and B) H&E-stained images of aortic root sections derived from 14-wks old Western diet-fed smOGT^WT^;ApoE^-/-^ and smOGT^KO^;ApoE^-/-^ mice. Corresponding summary graphs for lipid burden and plaque area are shown in C) and D).

### SMC-specific OGT deficiency abrogates inflammatory and smooth muscle cell lesion abundance in western diet-fed hyperglycemic ApoE^-/-^ mice

Aortic root lesions formed in Western diet-fed hyperglycemic smOGT^WT^ApoE^-/-^ mice (with intact OGT) revealed remarkable leukocyte infiltration and SMC content, as shown via immunohistochemistry. In contrast, SMC-specific OGT ablation reduced both leukocyte and SMC abundance within aortic root lesions of Western diet-fed smOGT^KO^ApoE^-/-^ mice (**Figures 8A, 8B**). Specifically, CD45 expression depicting leukocyte content was significantly diminished in the aortic root lesions of smOGT^KO^ApoE^-/-^ mice lacking smooth muscle OGT reflecting reduced inflammatory lesion burden in these animals (p<0.005 vs. smOGT^WT^ApoE^-/-^; **Figure 8C**). Similarly, immunofluorescence quantification revealed a significant decrease in ACTA2-positive area in aortic root sections derived from western diet-fed smOGT^KO^ApoE^-/-^ genotypes, illustrating attenuated SMC abundance in these animals (p<0.0001 vs. smOGT^WT^ApoE^-/-^; **Figure 8D**). In each case, comparable tissue sections incubated with an isotype-specific IgG antibody (instead of the corresponding species-specific primary antibody) displayed negligible background staining, confirming staining specificity for each antibody. Together, these results clearly demonstrate that SMC-specific lack of OGT blocks inflammatory and SMC lesion invasion in hyperglycemic ApoE^-/-^ mice *in vivo*.

**Figure 8.**
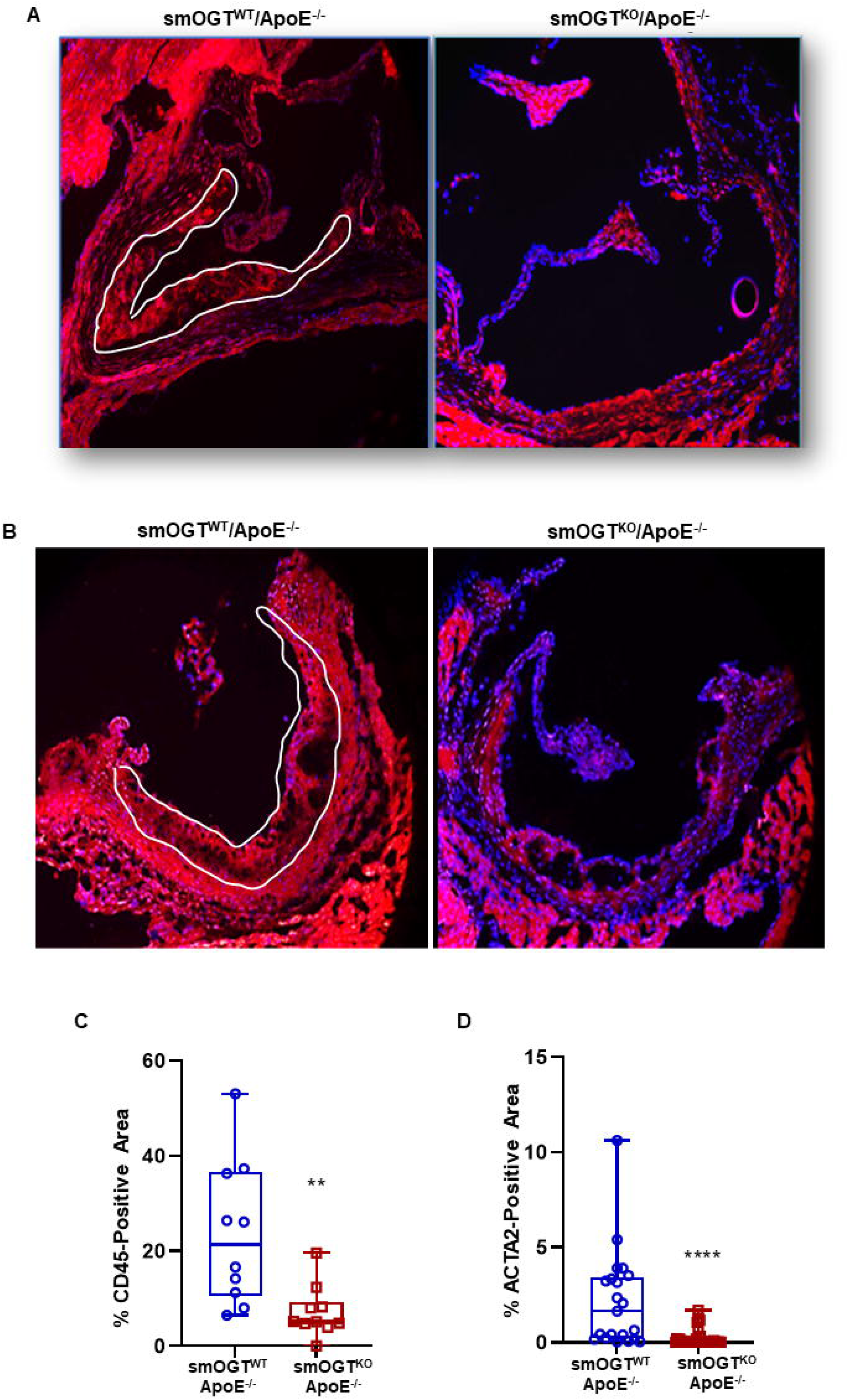
SMC-specific OGT deletion reduces inflammatory and smooth muscle cell content in western diet-fed ApoE^-/-^ mice. Shown are representative immunofluorescent images depicting A) CD45 and B) ACTA2 expression in aortic root sections derived from Western diet-fed smOGT^WT^;ApoE^-/-^ and smOGT^KO^;ApoE^-/-^ mice. Corresponding summary graphs are shown in C-D.

### Lack of SMC-specific OGT increases SM contractile marker expression while reducing inflammatory and proliferative marker expression in Western diet-fed ApoE^-/-^ mice

To delineate the role of smooth muscle OGT in VSMC differentiation and inflammatory response, aortic lysates from Western diet-fed smOGT^KO^ApoE^-/-^ and smOGT^WT^ApoE^-/-^ mice were subjected to immunoblotting. As shown in **Figure 9**, SMC-targeted OGT deletion significantly augmented LMOD1 (2-folds; p<0.005) and ACTA2 (3-folds; p<0.05) expression (SM contractile markers) in aortic vessels of western diet-fed smOGT^KO^ApoE^-/-^ mice vs. smOGT^WT^/ApoE^-/-^ mice with intact OGT. Under similar experimental conditions, loss of SMC-specific OGT reduced PCNA expression (proliferation marker) accompanied with diminished ERK phosphorylation in smOGT^KO^ApoE^-/-^ mice (p<0.05). Real-time PCR experiments further revealed that SMC-specific OGT deficiency significantly augmented the relative mRNA expression of classical SM contractile genes in western diet-fed smOGT^KO^ApoE^-/-^ aortic vessels compared to age-matched smOGT^WT^ApoE^-/-^ mice (**Figure 9H**). Specifically, both smooth muscle actin (*Acta2*) and calponin (*Cnn1*) mRNA levels were elevated in ApoE^-/-^ mice lacking SMC-specific OGT (p<0.05 vs. ApoE^-/-^ with intact OGT). Under similar experimental conditions, increased SM contractile mRNA expression was accompanied with downregulation of *Il6* and *Il1*β mRNA levels in mice lacking smooth muscle OGT. Specifically, both *Il6* and *Il1*β mRNA levels were significantly attenuated in Western diet-fed smOGT^KO^ApoE^-/-^ vs. ApoE^-/-^ mice with intact OGT (**Figure 9G**). Together, these data clearly demonstrate that SMC-specific OGT deletion promotes SMC differentiation in aortic vessels of hyperglycemic ApoE^-/-^ mice evidenced via upregulated SM contractile and reduced inflammatory and proliferative marker expression.

**Figure 9.**
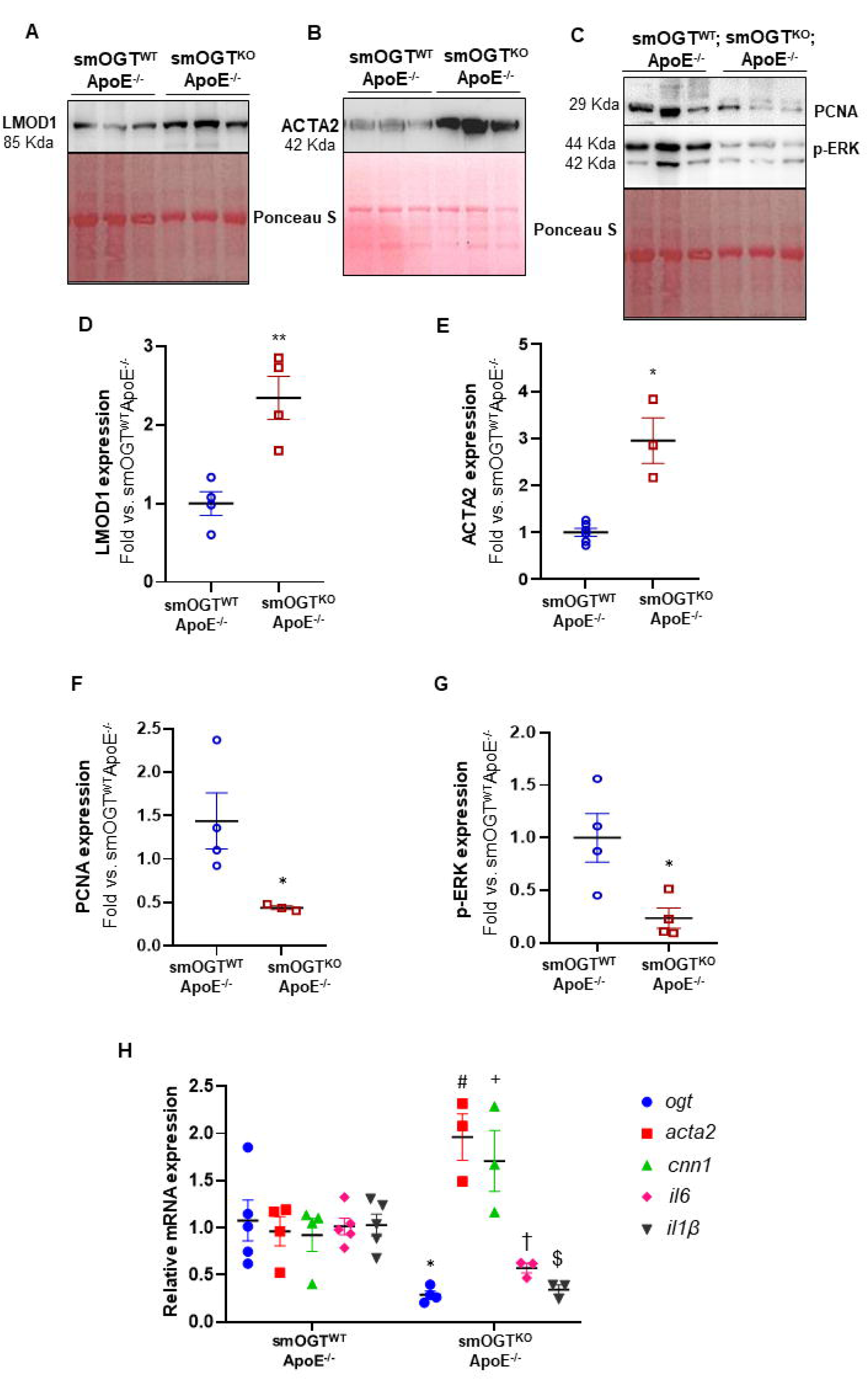
SMC-specific OGT deletion increases SM contractile marker expression concomitant to reduced proliferative and inflammatory marker expression in aortic vessels of western diet-fed ApoE^-/-^ mice. Aortic lysates derived from western diet-fed smOGT^WT^;ApoE^-/-^ and smOGT^KO^;ApoE^-/-^ mice were subjected to immunoblotting. Shown are A-C) representative immunoblots and D-G) corresponding summary graphs depicting LMOD1, ACTA2, PCNA and pERK expression. H) relative mRNA expression of *Ogt*, *Acta2*, *Cnn1*, *Il6* and *Il1*β.

### SMC-specific OGT deletion attenuates YY1 and SRF expression in aortic vessels of hyperglycemic mice *in vivo*

Concomitant to increased SM contractile marker expression, immunoblotting of aortic lysates prepared from smOGT^KO^ApoE^-/-^ mice revealed attenuated YY1 and SRF expression (transcriptional regulators of SM contractile genes). Specifically, YY1 and SRF protein expression were significantly diminished in the vascular walls of western diet-fed ApoE^-/-^ mice with SMC-specific OGT deletion (4-6-fold vs. smOGT^WT^ApoE^-/-^, p<0.0001; **Figures 10A, 10B**). Consistently, both SRF and YY1 protein expression were markedly reduced in the aortic vessels of STZ-induced hyperglycemic smOGT^KO^ mice. Specifically, STZ-induced hyperglycemia upregulated YY1 expression in smOGT^WT^ aortic vessels (p<0.0005 vs. normoglycemic smOGT^WT^). In contrast, lack of SMC-specific OGT profoundly diminished YY1 expression in hyperglycemic smOGT^KO^ mice (4-fold vs. hyperglycemic smOGT^WT^; p<0.0001, **Figure 10D**). Similar to these results, SMC-specific OGT deficiency downregulated SRF expression in smOGT^KO^ mice compared with smOGT^WT^ with intact OGT, and this effect was observed under both basal and hyperglycemic conditions (3-4-fold, p<0.0005, **Figure 10C**). Interestingly, while hyperglycemia increased YY1 expression in the vascular walls of smOGT^WT^ mice, SRF expression was significantly elevated in smOGT^WT^ mice both basally and in response to STZ-induced hyperglycemia, with no significant increase between these animals. Finally, as shown via real-time PCR, *Myocd* mRNA expression was upregulated in aortic tissues of STZ-induced hyperglycemic smOGT^KO^ mice lacking OGT compared with hyperglycemic smOGT^WT^ genotypes with intact OGT (p<0.05; **Figure 10E**). Collectively, these results demonstrate that smooth muscle OGT modulates YY1 and SRF expression under hyperglycemic conditions.

**Figure 10.**
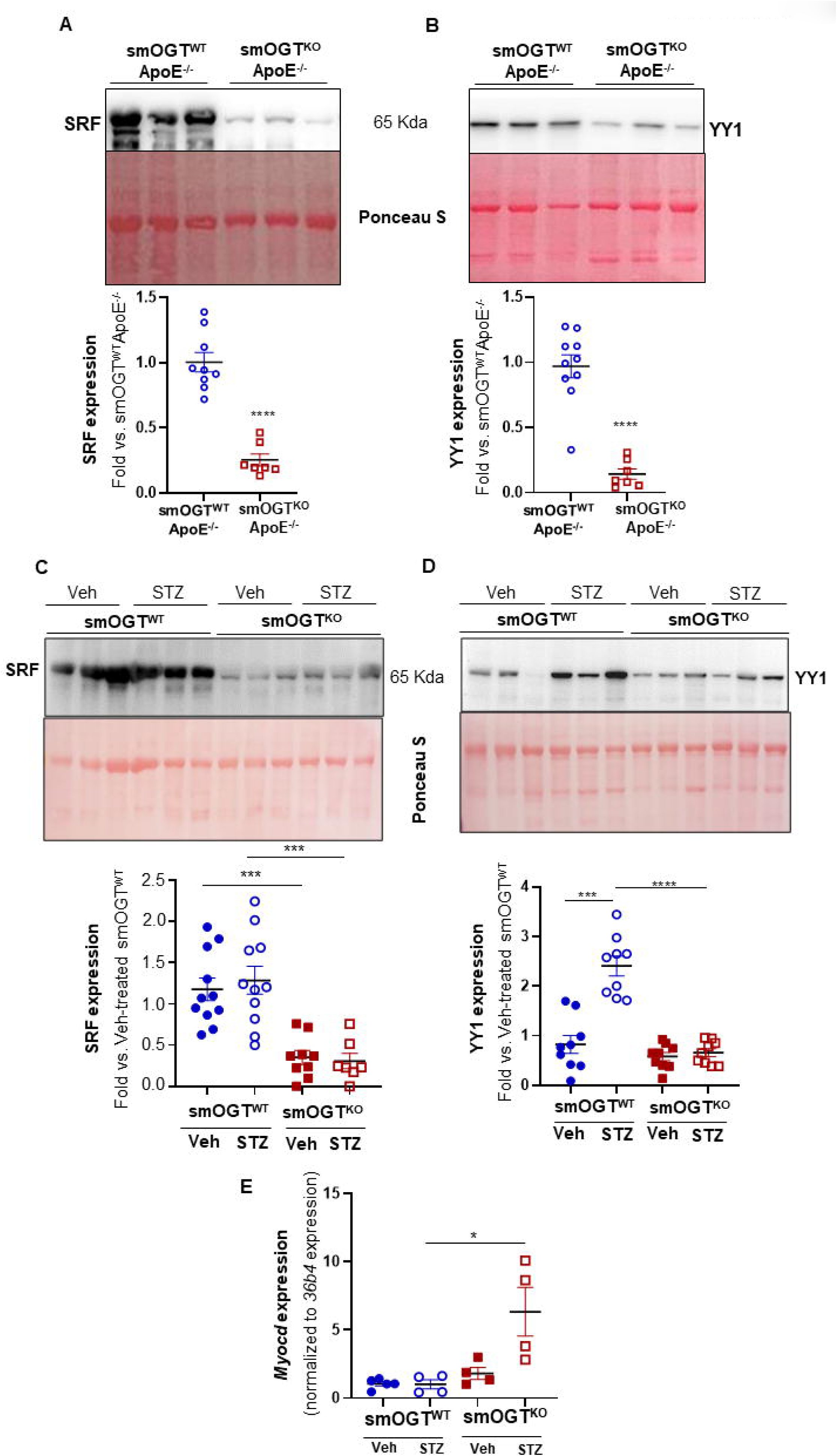
SMC-specific OGT deletion attenuates YY1 and SRF expression in aortic vasculature of western diet-fed ApoE^-/-^ and STZ-induced hyperglycemic mice *in vivo*.

## Discussion

The present study provides novel evidence for a causal role of smooth muscle OGT and O-GlcNAcylation in hyperglycemia-induced atherosclerosis. Although earlier work has reported that OGT-mediated O-GlcNAcylation promotes diabetic vascular calcification^25^, the conceptual link between SMC-derived OGT and hyperglycemia-driven lesion pathogenesis has not been previously explored. In this report, we provide the first demonstration that smooth muscle OGT-dependent mechanism(s) drive atherosclerotic lesion formation stimulated by hyperglycemia. Notably, our data suggests that OGT-mediated O-GlcNAcylation regulates VSMC de-differentiation in response to hyperglycemia.

Previous studies have reported that activation of the hexosamine biosynthetic pathway and O-GlcNAc signaling are key nutrient sensors modulating leptin expression in adipose tissues. OGT signaling has been shown to play a role in adipose tissue homeostasis. Specifically, OGT was reported to inhibit visceral fat lipolysis, with adipocyte-derived OGT mediating brain-to-adipose signaling prompting hyperphagia and diet-induced obesity. Moreover, absence of OGT in adipocytes was found to prevent diet-induced insulin resistance coupled with reduced circulating concentrations of free fatty acids^38, 39^. In the current work, loss of SMC-specific O-GlcNAc signaling did not affect either body weight, glycemic index, nor lipid levels in the hyperglycemic mice *in vivo*. Metabolic parameters including maximal oxygen consumption (VO_2_), carbon dioxide production (VCO_2_), physical activity, food intake, and energy expenditure also remained unaffected in mice lacking smooth muscle OGT, and this effect was observed under both hyperglycemic and non-hyperglycemic conditions. These results concur with an earlier report showing comparable body weight, oxygen consumption, food intake, and physical activity in liver-specific OGT knockout mice compared to wild type animals with intact OGT signaling^40^. Cogent to these earlier studies, the lack of difference between the metabolic phenotype of wild type vs. SMC-specific OGT knockout mice observed in the current study reinforces the contribution of adipose tissue O-GlcNAc signaling as a fat-sensing PTM.

Growing literature indicates that O-GlcNAcylation affects multiple physiological and pathophysiological processes relevant to the cardiovascular system. It is well established that protein O-GlcNAcylation can affect numerous cellular functions via regulation of transcription, metabolism, cellular localization, mitochondrial function, protein turnover and protein stability as well as autophagy. Previous studies have shown that increased O-GlcNAcylation of cardiac proteins may contribute to the adverse effects of diabetes in the heart and cardiovascular system. Specifically, OGT ablation in cardiomyocytes was reported to cause dilated cardiomyopathy with systolic and diastolic dysfunction^26, 41^ Conversely, recent studies using in vitro and in vivo murine models of ischemia-reperfusion injury have demonstrated cardioprotective responses of elevated protein O-GlcNAcylation^42^. In an inducible cardiomyocyte-specific OGT knockout murine model, it was reported that conditional OGT ablation in cardiomyocytes leads to progressive ventricular dysfunction in the adult mice, supporting a critical role of cardiomyocyte OGT in mammalian heart maturation^43^. In the current study, inducible SMC-specific OGT deletion did not result in an adverse cardiac phenotype. This data validates the smooth muscle specificity of the Cre murine driver *Myh11-CreER^T^*^2^ employed, devoid of any off-target effects.

Clinical and animal data, including genetic association studies, have linked hyperglycemia, a hallmark feature of Type 1 and Type 2 diabetes, with increased O-GlcNAc signaling^19, 44, 45^. Extensive literature, including our earlier work, indicates that augmented protein O-GlcNAcylation is a putative trigger for diabetes-related complications. We previously reported increased atherosclerotic lesion formation in hyperglycemic ApoE^-/-^ mice accompanied with elevated OGT and O-GlcNAc protein expression in the vasculature^34^. Recent studies have shown that OGT-mediated O-GlcNAcylation modulates vascular calcification and delayed wound healing in diabetes. Congruent to these findings, in the present study we have demonstrated that SMC-specific loss of OGT resulting in reduced O-GlcNAcylation prevents atherosclerotic lesion formation in ApoE^-/-^ mice induced by Western diet feeding. It is important to note that western diet feeding increased fasting blood glucose levels in the ApoE^-/-^ mice, triggering a diabetic phenotype in these animals. Such an increase in glycemic levels was accompanied by augmented protein O-GlcNAcylation in the vasculature (**Figure 5**). These data are in accordance with earlier reports revealing elevated fasting glucose and insulin levels in ApoE^-/-^ mice following western diet feeding regimen^9, 10^. Importantly, we show that SMC-specific OGT loss impedes lipid burden and plaque area in hyperglycemic ApoE^-/-^ mice, in the absence of significant changes in blood glucose and plasma lipid levels compared with hyperglycemic ApoE^-/-^ mice with intact OGT. These results postulate a direct regulatory role of OGT-mediated O-GlcNAcylation in atherosclerotic lesion pathogenesis, independent of glucose or lipid alterations.

Several lineage tracing studies have established the contribution of vascular smooth muscle cells to atherosclerotic lesion formation^46–48^. De-differentiation of vascular smooth muscle cells is characterized by enhanced migratory, secretory and proliferative properties concomitant to loss of the contractile gene expression. Diabetic patients have an increased propensity for VSMC migration and proliferation, features characteristic of VSMC de-differentiation from a ‘quiescent’ contractile state to a ‘synthetic’ proliferative phenotype. We previously demonstrated that pharmacological activators of O-GlcNAc stoichiometry stimulate proliferation of human aortic smooth muscle cell (HASMC) primary cultures incubated with high glucose *in vitro*, reflective of the diabetic milieu. Conversely, pharmacological OGT inhibitors blocked glucose-stimulated HASMC proliferation^30^. We have further shown that augmented OGT and O-GlcNAc protein expression in aortic vessels of hyperglycemic ApoE^-/-^ mice associate with robust atherosclerotic lesion formation and enhanced proliferative VSMC lesion abundance^34^. More recently, a positive correlation was also reported between SMC de-differentiation and OGT-mediated O-GlcNAcylation in PDGF- and rapamycin-treated coronary SMC^49^. Further, both OGT and O-GlcNAc protein expression were augmented in a femoral artery endothelial denudation injury mouse model. Likewise, in a murine model of carotid artery ligation^50^, it was shown that while vascular injury increased OGT and O-GlcNAc protein expression, SMC-specific OGT loss-of-function reduced neointima formation. Consistent with these earlier reports, in the current work we have shown that SMC-specific lack of OGT resulting in reduced protein O-GlcNAcylation upregulated the expression of multiple SM contractile markers in the aortic vasculature of both STZ-induced hyperglycemic smOGT^KO^ and Western diet-fed hyperglycemic smOGT^KO^;ApoE^-/-^ mice. Our data lends strong support to the notion that OGT promotes VSMC de-differentiation in response to hyperglycemia. Characterization of lesion complexity further demonstrated an attenuation in SMC abundance and leukocyte infiltration in aortic root lesion area of hyperglycemic smOGT^KO^;ApoE^-/-^ mice vs. hyperglycemic smOGT^WT^; ApoE^-/-^ animals. This was further accompanied by downregulation of the proliferation marker PCNA expression in hyperglycemic ApoE^-/-^ with SMC-targeted OGT deletion. Our data that OGT gene silencing attenuates cellular proliferation is congruent to earlier reports^93–95^. For instance, Xu et al (2019) previously showed that cellular proliferation was markedly inhibited by OGT knockdown in colon cancer cell lines.^94^ Likewise, in pulmonary artery smooth muscle cells isolated from idiopathic pulmonary arterial hypertension lung explants, OGT gene silencing was found to reduce cell proliferation as revealed via BrdU staining.^95^ In the context of these earlier reports, our findings implicate a beneficial impact of SMC-targeted ablation of OGT signaling in hyperglycemia-induced VSMC proatherogenic responses. Collectively, the current work underscores a direct regulatory role of SMC-derived OGT in lesion pathogenesis in diabetes.

Extracellular signal-regulated kinase 1/2 (ERK1/2), a member of the mitogen-activated protein kinase (MAPK) family, participates in many cellular programs including cell proliferation, differentiation, motility, and cell death. There is accumulating evidence emphasizing a potential crosstalk between O-GlcNAcylation and ERK1/2 signaling pathways. HBP activation and elevated O-GlcNAc levels have been reported to stimulate ERK1/2 and p38-MAPK phosphorylation. Earlier studies have also suggested a role of MAPK signaling in regulation of VSMC phenotypic transition, attributing to SMC heterogeneity and vascular pathology. In accordance with these earlier findings, we observed a significant reduction in pERK expression, a key signaling mediator of VSMC growth, in aortic lysates derived from smOGT^KO^;ApoE^-/-^ vs. smOGT^WT^;ApoE^-/-^ mice with intact OGT following Western diet-induced hyperglycemia. Concomitant to reduced ERK phosphorylation, SMC-targeted OGT deletion attenuated proinflammatory markers (IL1β and IL6) typically elevated in diabetic vascular pathology. Contrary to these results, an earlier work by Zhao et. al. (2018) has revealed increased intestinal epithelial hypertrophy, epithelial hyperplasia, and mucosal thickness in intestinal epithelial-specific OGT knockout (Vil-OgtKO) mice, coupled with increased inflammatory gene expression (Tnfα, IL6, IL1β). The observed discrepancy in these results may be due to cell-specific responses of OGT gene silencing in two diverse cell types (SMC vs. epithelial cells), prompting disparate pathologies.

Regulation of VSMC differentiation is a critical event through embryogenesis and plays a key role during vascular remodeling in response to proatherogenic stimuli (such as hyperglycemia) The fate of de-differentiated VSMC in atherosclerosis involves a transitional, multipotent state with context-dependent adoption to other cell types facilitating disease progression^14^. Early literature suggests that VSMC phenotypic modulation is a function of numerous factors including inflammatory mediators, cytokines and growth factors, and extracellular matrix proteins^51, 52^. Increasing evidence further demonstrates that a triad of transcription factors (TFs) and co-regulators mediate the transcriptional regulation of de-differentiated VSMC gene expression while disrupting the normal genetic program for VSMC contractile function. Interaction of one such TF called the serum response factor (SRF) with its co-factor Myocardin (MYOCD) is recognized as the central regulator of VSMC contractile gene transcription^53^. In contrast, multiple transcriptional repressors including YY1, Elk-1 and KLF4 pathways block SRF-MYOCD interaction, repressing the SMC contractile phenotype^54–56^. Despite extensive efforts, significant gaps persist in our understanding of how diabetes regulates the molecular pathways mediating VSMC phenotypic transition. Growing data indicates that O-GlcNAcylation of transcriptional proteins modulates numerous cell functions relevant to diverse pathological conditions. In a recent study, O-GlcNAc-mediated activation of RUNX2, an osteogenic TF, was reported to modulate VSMC transdifferentiation to osteoblast-like cells, in turn promoting diabetic vascular calcification^25, 57^. Findings from the present study prompt us to speculate that YY1- and SRF-dependent pathways play a regulatory role in OGT-mediated SMC de-differentiation induced by hyperglycemia, promoting accelerated atherosclerotic lesion formation in diabetes. Recruitment of SRF-MYOCD complex on the CArG box, a highly conserved cis-regulatory element (CC(A/T)6GG) found in the promoters of most SMC-specific genes, stimulates multiple SMC contractile gene transcription (*Sm22*α, *Acta2*, S*m-mhc*)^15, 16^.^15^. Growing literature indicates that the nuclear protein SRF can mediate disparate target gene expression, stemming from its ability to associate with alternating classes of co-factors^58, 59^. Consequently, a dual SRF function in VSMC phenotypic regulation is likely contingent upon its binding partners, with SRF-Myocd complex mediating SMC differentiation, while interplay of SRF with ternary complex factors (TCF), including ETS-like proteins (ELK) and Ying yang 1 (YY1), controlling cell cycle entry^55, 58, 60^. While the latter regulates expression of many immediate early genes (c-fos, Egr-1), the former is known to control SMC differentiation genes^58–61^. Indeed, an earlier study identified SRF as a key regulator of VSMC proliferation and senescence. Our data are comparable to these previous findings. Specifically, we observed that SMC-specific OGT deletion reduced SRF expression concomitant to attenuated PCNA expression in the aortic vasculature of smOGT^KO^;ApoE^-/-^ vs. smOGT^WT^;ApoE^-/-^ mice in response to Western diet-induced hyperglycemia. While these data reinforce the contribution of SRF in VSMC proliferation, our findings suggest a novel regulatory role of OGT-mediated O-GlcNAcylation in this process. YY1, on the other hand, is a ubiquitous multifunctional TF belonging to the GLI-Krüppel family of zinc-finger proteins. It is known to be involved in embryonic development,^119^ and a number of pathological processes such as cytokinesis, apoptosis, cell differentiation, and cell cycle.^115^ Multiple studies indicate that YY1 can facilitate both transcriptional gene activation and gene repression in turn regulating numerous cellular processes including cell growth, cell division, differentiation and replication^62–64^. For instance, YY1 was reported to bind to the CArG box in the SM22 promoter and operate as a transcriptional activator in SMCs;^19;^ on the other hand, YY1 has also been shown to repress smooth muscle promoter activity by competitively displacing MYOCD from SRF inhibiting the MYOCD/SRF/CArG box-mediated gene transcription.^20, 21^ ^65, 66^. Additional work has demonstrated that YY1 controls gene transcription either by directly binding to the promoter or acting as a cofactor of several transcription factors.^18^ Cogent to its dual role in transcriptional regulation, it was hypothesized that YY1 facilitates cell cycle regulation via cell-specific mechanisms^64, 67–69^. An earlier study demonstrated increased YY1 expression in vascular SMC in response to mechanical vascular damage, thereby blocking SMC replication *in vitro*.^120^ More recently, Zheng et al. (2020) have reported that vascular injury induces YY1 expression in carotid artery medial SMCs in association with reduced transcription of SMC differentiation marker.^109^ YY1 overexpression has also been shown to decrease VSMC cell growth and migration.^121^ Accordingly, existing literature supports an inhibitory role of YY1 on SMC differentiation, possibly acting as a transcriptional repressor. Our findings that hyperglycemia increased YY1 expression in aortic vessels of smOGT^WT^ mice while SMC-specific OGT deletion reduced YY1 levels concomitant to augmented SMC contractile marker and attenuated cell proliferation marker expression are in accordance with these earlier reports. As such, we speculate that interaction of YY1- and SRF-dependent pathways drive OGT-mediated SMC transformation to a de-differentiated phenotype in response to hyperglycemia. While our data suggests a putative crosstalk between YY1/SRF-related mechanisms and OGT-mediated VSMC de-differentiation, additional studies are warranted to elucidate the molecular pathways that link OGT to YY1 and SRF in regulation of VSMC phenotypic transformation in diabetes. We further posit that OGT-mediated O-GlcNAcylation of YY1 regulates SRF-dependent pathways and binding partners, in turn advancing VSMC cell cycle progression, VSMC-to-macrophage fate switch and inflammatory VSMC phenotype in diabetes.

A limitation of this study relates to the y-linked Cre murine driver employed that restricts interrogation of sex-based differences in OGT-mediated SMC de-differentiation. Given that SMCs play a prominent role in vascular pathology, and vascular disease accounts for multiple cardiovascular anomalies striking men and women alike, it would be important to decode the molecular link between OGT and diabetic vasculopathy across both sexes. To this end, future investigation currently underway in our lab will utilize a VSMC-restricted Cre driver mouse (*<u>Itga8-CreER^T^</u>*^2^) to enable sex-specific studies and to selectively knockout OGT in VSMCs of vessel wall while sparing visceral SMC^70^. Notably, using VSMC-specific reporter mice combined with VSMC-specific OGT deletion, additional work will interrogate the spatiotemporal contribution of OGT-mediated O-GlcNAcylation in VSMC fate switch to diseased phenotypes and delineate underlying molecular mechanisms.

In conclusion, the present study uncovers a protective outcome of SMC-specific OGT deletion in hyperglycemia-induced atherosclerosis. Our data underscores a conceptual link between OGT-mediated O-GlcNAcylation and SMC de-differentiation and highlights smooth muscle OGT as a putative target of hyperglycemia-induced vasculopathy. Future work will determine whether OGT-mediated O-GlcNAcylation drives VSMC fate transition during lesion pathogenesis in diabetes. In addition, elucidation of the temporal course and contribution of smooth muscle OGT to lesion regression in diabetes is warranted.

